# Experimental infection characteristics of *Bordetella* pertussis via aerosol challenge on rhesus macaques

**DOI:** 10.1101/356022

**Authors:** Dachao Mou, Peng Luo, Jiangli Liang, Qiuyan Ji, Lichan Wang, Na Gao, Qin Gu, Chen Wei, Yan Ma, Jingyan Li, Shuyuan Liu, Li Shi, Mingbo Sun

**Affiliations:** Institute of Medical Biology; Chinese Academy of Medical Science & Peking Union Medical College, Kunming, Yunnan, China.; Yunnan Key Laboratory of Vaccine Research and Development on Severe Infections Diseases, Kunming, Yunnan, China; Department of Diphtheria, Tetanus and Pertussis vaccine and Toxins, National Institute for Food and Drug Control, Beijing, China

**Keywords:** Pertussis, Rhesus macaque animal model, Aerosol challenge

## Abstract

The effect of aerosol challenge of rhesus macaques with *Bordetella pertussis* and the feasibility of using rhesus monkeys as an animal model for pertussis infection were evaluated in this study. Four 1-year old rhesus macaques were aerosol challenged with *B. pertussis* at the concentration of 10^5^ CFU/mL for 30 min (group 1) or 60 min (group 2). Rectal temperature was found slightly increased at days 3 and 5 and returned to baseline levels at day 21 after challenge. White blood cell counts peaked at day 7, with a 4.7~6.1-fold increase and returned to baseline levels at day 45. Bacteria colonization of nasopharyngeal swabs was observed, and the number of colonies was gradually increased and peaked at day 14, reaching 5.4-8.1 × 10^6^/mL. The seroconversion rate of anti-pertussis toxin (PT), pertactin (PRN), and filamentous hemagglutinin(FHA) antibodies was 100%, with an increase in geometric mean titers after challenge. Analysis of cytokines revealed that the levels of cytokines including IL-2, IL-6, IL-8, IL-10, IL-17A, IL-13, IL-12, and IL-18 were significantly increased at days 5 to 14 in group 2. These results demonstrate that the characteristic of pertussis infection in infant rhesus macaque was similar as in human beings, which provide a clue to using infant rhesus macaque as a candidate model for pertussis infection in the future studies

## Introduction

Pertussis is a contagious respiratory disease caused by the bacterium Bordetella pertussis. The disease is characterized by sudden spasmodic cough accompanied by inspiratory whoop or posttussive emesis lasting several months and may cause death in infants without treatment[1]. Guillaume De Baillou first described a pertussis outbreak in Paris in the summer of 1578[2]. The epidemic involved mainly infants, and the mortality rate was high. The diagnosis of pertussis was based primarily on bacterial culture[3, 4], antigen detection by polymerase chain reaction[5], and serological antibody detection[6]. Pertussis remains a major threat to the health of children worldwide and causes high economic burden at the state and local level[7, 8].

Pertussis is preventable and, in the 1930s, Medson [9] developed whole-cell pertussis (wP) vaccine. In 1981, Japanese researchers developed acellular pertussis (aP) vaccine containing pertussis toxin (PT) and filamentous hemagglutinin (FHA) or a small amount of agglutinogen[10, 11]. The vaccine significantly reduced the incidence of pertussis. However, in recent years, the effectiveness of pertussis vaccine has been contested. In the United States, although the pertussis vaccination rate is >95%, the incidence of pertussis in the past 30 years is increasing[12], especially in 2004, 2010, and 2012[13, 14]. This increase may be related to the decrease in the immune protective efficiency induced by vaccination[15], transmission of asymptomatic latent infection[16], cross-infection via healthcare personnel and travelers[17, 18], and pathogen adaptation in immune populations after large-scale vaccination [19].

The immune mechanisms underlying pertussis infection and the efficacy of new vaccines and combined vaccines can be studied by establishing suitable animal models that replicate the full spectrum of the disease. Most infection studies are carried out in mouse, rat, rabbit, and piglet models of pertussis[20, 21]. However, these models cannot reproduce the full clinical spectrum achieved in human models[22, 23]. With respect to studies using non-human primate models, a successful baboon model was established by Warfel et al[24]. However, the use of baboons as an animal model of pertussis is limited because of the limited availability of animals, high housing costs, and lack of suitable reagents. Compared with the baboon model, the advantages of using rhesus macaque models include high availability of animals, low housing costs, and availability of suitable reagents, and therefore this animal model is considered ideal[25]. Since 1929, three studies have evaluated the infection of rhesus monkeys with *B. pertussis*. However, these studies could not replicate the clinical disease in humans[26–28].

In the present study, rhesus macaques aged 12 to 14 months were challenged with *B. pertussis* via aerosol injection; indices of infection, including leukocyte count and colonization of the nasopharynx, were determined; and the humoral immune response and cytokines were analyzed to assess the possibility of using rhesus monkeys as an animal model for pertussis infection. Infection efficiency was evaluated after challenge for different periods.

## 1 Materials and methods

### 1.1 *B. pertussis* medium preparation

A total of 44 g of Bordet-Gengou agar medium (B-G medium: peptone, 20.0 g/L; potato extract, 4.5 g/L; NaCl, 5.5 g/L; agar, 14.0 g/L, pH 6.7 ± 0.2 from Haibo Biology) and 10 mL of glycerol were added to 700 mL of distilled water. The medium was mixed, dissolved by heating, and autoclaved at 121 °C for 15 min. The solution was cooled to 45–50 °C, and 300 mL of sterile defibrinated sheep blood (Lanzhou Minhai Company) was added. The culture medium was transferred to a sterile Petri dish or test tubes. *B. pertussis* (strain No.18323/CMCC58030; batch No. 2012003, obtained from the National Institutes for Food and Drug Control and cryopreserved by IMBCAMS) was resuspended in a small volume of sterile saline and transferred to a flask containing culture medium. The medium was incubated in an incubator at 37 °C and transferred to two flasks containing culture medium. After incubation at 37 °C for 48 h, the medium was transferred to four to six flasks and incubated at 37 °C for 24 h. The bacterial sediment was resuspended in isotonic saline, diluted to a concentration of 10^11^ CFU/mL using a turbidimetric method, and used within 2 h after preparation.

### 1.2 Animal use and welfare

Four healthy rhesus macaques, two males and two females, aged 12 to 14 months, were obtained from the Institute of Medical Biology, Chinese Academy of Medical Sciences (IMBCAMS) (Animal License No. SCXK (Dian) K2015-0004) The rhesus macaques were assigned to two groups randomly, with two animals in each group (one male and one female) Animal study was conducted in compliance with the Animal Welfare Act, Declaration of Helsinki (2013 revision) and other regulations relating to animal experiments. The rhesus macaques were confirmed to be healthy, acclimated to the animal house, and identified by hanging tag on the neck And these animals were fed with food and fruits strictly complying with requirements of animal welfare and music was played during 11:00 ~13:00 every day.

### 1.3 Challenge of rhesus macaques

The aerosol apparatus used for this study was designed by our lab and produced by Lanfang Honlan Equipment Co. The apparatus was composed of a rectangular Plexiglas chamber with a removable lid (40 cm × 60 cm × 40 cm), a pump and a medical nebulizer (average atomization rate: ≥ 0.15 mL/min, working pressure: 60–150 KPa, normal working condition: 10–40 °C). The pump was connected to the inlet side of the nebulizer to deliver *B. pertussis* suspension for atomization. The outlet side of the nebulizer was connected to two inlet port of the challenge chamber to deliver atomized *B. pertussis* to the interior of the chamber. Outlet tube with an air filter was connected to the challenge chamber to remove air. An air sampling port was embedded in the middle of the challenge chamber to monitor actual concentration of aerosol *B. pertussis* inside the chamber(Fig 1).

**Fig. 1.**
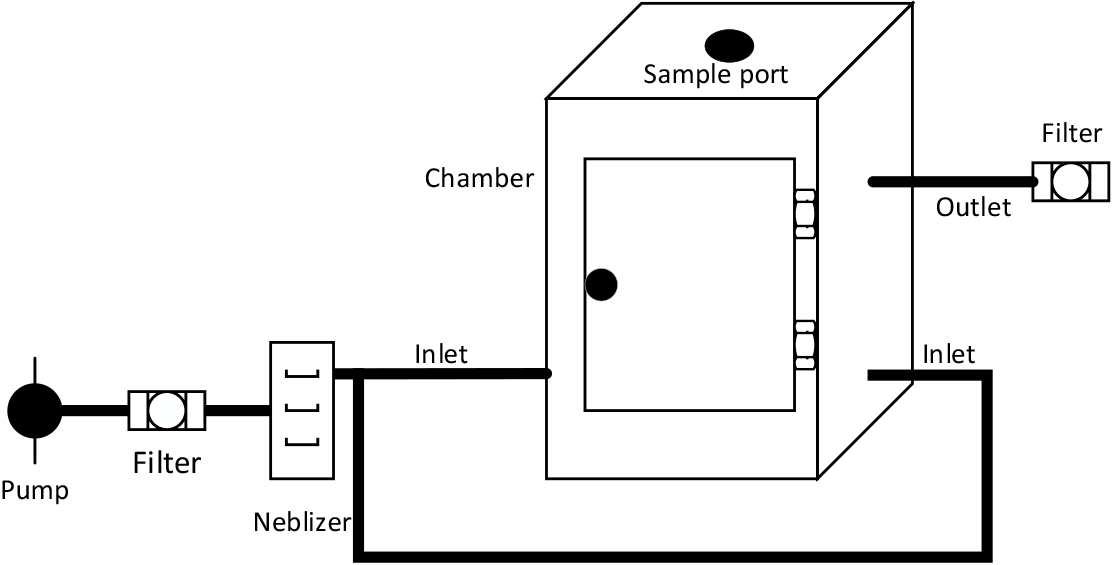
Simplified layout of aerosol apparatus

*B. pertussis* strain 18323 at a concentration of 10^11^ CFU/mL with 8mL was delivered to the nebulizer for aerosolisation. After aerosol generation, 10 mL of air was sucked out from the sampling port every 5 min using syringes and injected into 10 mL saline. The solution was transferred to B-G medium and the number of colonies was determined to ensure actual challenge concentration.

The rhesus macaques of one group were placed into the challenge box for challenging. In group 1, the macaques were numbered as 1 and 2, and were challenged with 10^5^ CFU/mL for 30 min. In group 2, the macaques were numbered as 3 and 4 and were challenged with 10^5^ CFU/mL for 60 min. After challenging, the two groups were housed separately.

### 1.4 Monitoring the rectal temperature of rhesus macaques

The rectal temperature in the two groups was measured before aerosol challenge and at days 3, 5, 7, 10, 14, 21, 31, and 45 after challenge.

### 1.5 White blood cell (WBC) count

A volume of 150 μL of EDTA-anticoagulated venous blood was drawn before aerosol challenge and at days 3, 5, 7, 10, 14, 21, 31, and 45 after challenge. The number of WBCs was determined by blood cell counting.

### 1.6 Culture of nasopharyngeal bacteria

At each sampling point, nasopharyngeal bacteria were collected using nasopharyngeal and nasal swabs. Bacteria were suspended in 1 mL of saline and cultured in B-G medium containing 30% sterile defibrinated sheep blood diluted 100, 400, 800, and 1600-fold within 2 h. The number of colonies was determined after 5 days. Colonies were picked from the blood agar, subjected to Gram staining, and their morphology was analyzed by microscopy. Bacterial DNA was extracted and identified by 16S rRNA sequencing. Sequencing was performed by Beijing Nuohezhiyuan Technology CO., Ltd. in an Illumina technology sequencing platform using the paired-end method.

### 1.7 Determination of antibodies of rhesus macaques after challenge

The venous blood of the two animal groups was drawn before aerosol challenge and at days 7, 10, 14, 21, 31, 45, and 60 after challenge. Serum was separated and anti-PT, PRN, FHA antibody titers were measured using ELISA. A 96-well microplate was coated with 3 μg/mL of antigens PT, PRN, and FHA and incubated at 4 °C overnight. Diluted serum was added to the microplate and incubated at 37 °C for 1 h (All the antigens are come from Department of DTP Vaccine and Toxin, National Institute for Food and Drug Control, China). After that, HRP-labeled sheep anti-monkey IgG was added to the microplate, and the plate was incubated at 37 °C for 1 h. The substrate 3, 3′, 5, 5′ tetramethylbenzidine (TMB) was added to the microplate, the plate was incubated at room temperature for 30 min, and the reaction was terminated by adding H_2_SO_4_. OD values were measured at 450 nm. A blank sample was included in each plate and OD values ≥ 2.1-fold those of the blank sample were considered positive. The antibody titers of different types were determined by calculating the geometric mean titer (GMT), as follows: GMT = Lg – 1 [(LgX1 + LgX2 +…LgXn)/n]. Seroconversion was defined as reaching a positive status (titer was higher than 1:200) after immunization in seronegative subjects (when the baseline titer was below 1:200) or as a 4-fold increase in titer after immunization when the baseline titer was higher than seronegative.

### 1.8 Measurement of cytokines after challenge

The venous blood of the two groups was drawn before aerosol challenge and at days 3, 5, 7, 10, 14, 21, 31, 45 and 60 after challenge. Serum was separated and 16 different cytokines—GM-CSF, G-CSF, IFN-, IL-2, IL-10, IL-15, IL-17A, IL-13, IL-1β, IL-4, IL-5, IL-6, TNFα, IL-12 (p40), IL-18, and IL-8—were tested using a Milliplex cytokine kit (Non-Human Primate Cytokines, Cat. No. PRCYTOMAG-40K, Merck Millipore, USA).

### 1.9 Statistics

The average values were calculated. Data were analyzed using a *t*-test and software SPSS version 22.0. P values of <0.05 were considered statistically significant. The fluorescence of cytokines was measured using a Luminex 200 system and analyzed in Milliplex Analyst software version 5.1.0.0 using the 5-PL method. Seven measurements were obtained for constructing the standard curve.

## 2 Results

### 2.1 Dose of aerosol used for challenge

Air sample was collected and tested by bacterial culture during aerosol challenge. Viable bacterial count of air samples 5 min after challenge was increased significantly, reached 10^5^ CFU/mL 10 min after challenge, and this level was maintained until the end of the study period (Fig. 2).

**Fig. 2.**
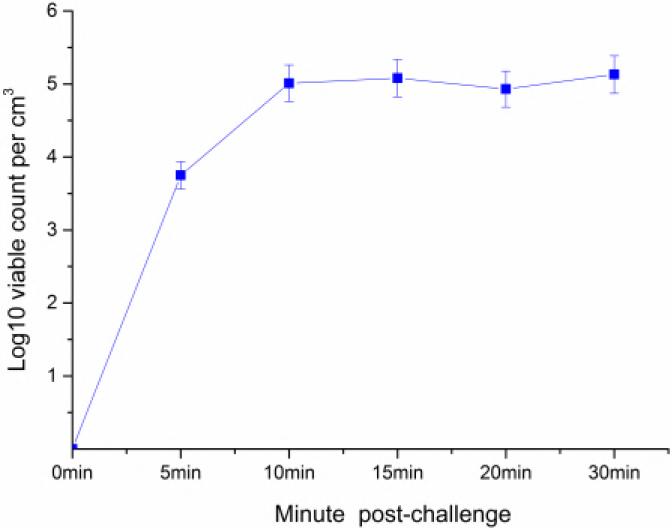
Viable count test for the air in the challenge box during aerosol challenging process. After aerosol challenge, 10 mL air was sucked out from sampling port using syringes 0, 5, 10, 15, 20, 30 min after challenge, and injected into 10mL saline, 100μL of the liquid was spread on the B-G medium containing 20% sheep blood and incubated at 37°C for 4 days. The number of colonies was determined.

### 2.2 Aerosol challenge with *B. pertussis* induced changes in vital signs and leukocytosis

Seven days after challenge with atomized *B. pertussis*, rhesus macaques developed pallor and cough, and coughing gradually disappeared after 21 days. However, as lack of suitable video recording system, coughing frequency and duration were not recorded. After challenge for 3 days, the rectal temperature of both groups was increased, peaked at days 5 and 7, and returned to normal levels at day 21 after challenge (Fig. 3A).

The number of WBCs in the two study groups was significantly increased at day 3 after challenge (P<0.05) and reach to the highest level at day 7 and returned to baseline values at day 45 (Fig. 3B). In group 2, WBC reached to 49,180 per μL, corresponding to a 6.1-fold increase relative to pre-challenge values, and decreased gradually. In group 1, WBC reached to 45,390 per μL, corresponding to a 4.7-fold increase, and decreased to 27310 per μL at day 10, then increased to 37330 per μL at day 14, and decreased gradually.

**Fig. 3 Dynamic profiles of rectal temperature and leukocytosis after challenged with *B.Pertussis*** Group 1 is challenged for 30 min and group 2 is challenged for 60 min with two rhesus macaques for each group. (A) Rectal temperature of two groups was measured at each time point before and after aerosol challenge with electronic thermometer. The data was presented as the average of the rectal temperature of each group, *p<0.05, +p<0.05. (B) The number of white bold cells was determined at each time point before and after aerosol challenging, *p<0.05, ***p<0.001.

### 2.3 Aerosol challenge increased nasopharyngeal colonization

At day 3 after challenge, swabs were collected from the nasal and nasopharyngeal of each test animal for bacterial culture. At days 5, 7, and 10 after challenge, the number of colonies was gradually increased and reached 5.4 × 10^6^ and 8.1 × 10^6^/mL (P<0.05) in groups 1 and 2, respectively. After 21 days, the number of colonies was decreased significantly (Fig. 4).

For bacterial identification, colonies collected from nasopharyngeal swabs (P1.5d, P2.5d, P3.5d, P4.5d) and nasal swabs (N1.5d, N2.5d, N3.5d, N4.5d) at day 5 after challenge and grown on Bordet-Gengou agar and *B. pertussis* grown on Bordet-Gengou agar were selected and identified by 16s rRNA sequencing. The results demonstrated that the sequences of colonies identified in nasopharyngeal swabs at day 5 after challenge were highly homologous to those of *B. pertussis* (35–95% as *B. pertussis).* In turn, the sequences of bacterial colonies identified in nasal swab samples at day 5 after challenge presented little homology to those of *B. pertussis* (the degree of homology to *B. pertussis* in one macaque was 25%). The bacteria were identified as Firmicutes and Actinobacteria, but not Proteobacteria. These results indicated that *B. pertussis* was present primarily in the nasopharyngeal (Fig. 5).

**Fig. 4.**
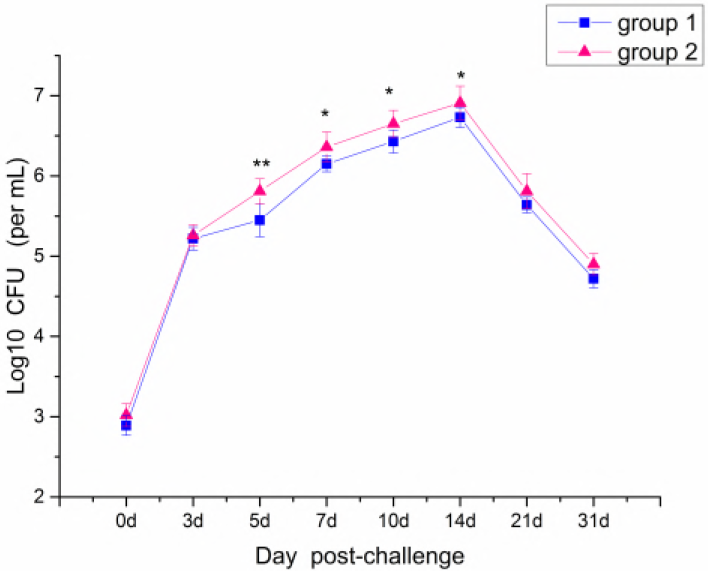
Colonization of nasopharynx of rhesus macaques after challenged with atomized *B.Pertussis*. At each sampling point, nasopharyngeal bacteria was taken using nasal and nasopharynx swabs, and suspended in 1mL saline. Similar as method described in Fig 2, 100μL was spread on cell culture, incubated at 37°C for 4 days and number of colonies was determined, *p<0.05, **p<0.01.

**Fig. 5.**
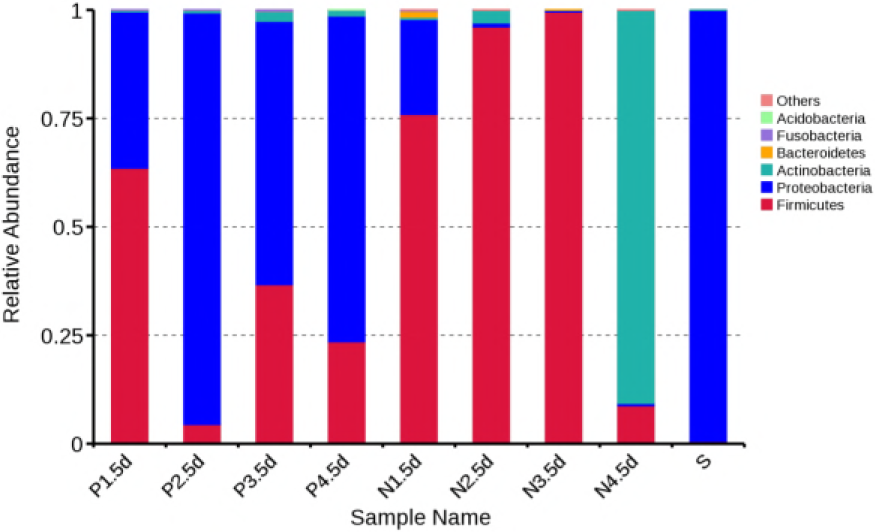
16s subunit identification result of colonies on blood plates. The suspected *B. Pertussis* colonies on the blood agar were picked and suspended in PBS. DNA of bacteria was extracted and identified using 16s subunit sequencing. P1-4 represented as bacteria recovered from **nasopharynx** swabs of No. 1 – 4 rhesus macaques, N1-4 represented as bacteria recovered from nasal swabs of No. 1 – 4 rhesus macaques and S serves as positive control of *B. Pertussis.*

### 2.4 Aerosol challenge induced specific antibody responses against *B. pertussis*

Anti-PT, PRN, FHA antibody titers were measured after challenge with B. pertussis. The seroconversion rate of antibody against PT reached 100% at day 14, and the seroconversion rate of antibody against FHA reached 100% at day 7 in both groups, the seroconversion rate of antibody against PRN reached 100% at day 7 and day 10 for group 1 and group 2, respectively. In group 1, the antibody level against PT, and FHA peaked after day 31 with GMTs of 4.41, 4.81, respectively, the antibody level against PRN peaked after day 14 with GMTs of 4.35, and declined afterward (Fig. 6A). In group 2, the antibodies induced by pertussis were significantly elevated, and the GMTs of the antibodies against PT, FHA, and PRN, were 4.81, 4.96, and 5.31, respectively. The titers tended to of decline, as in group 1 (Fig. 6B).

**Fig. 6.**
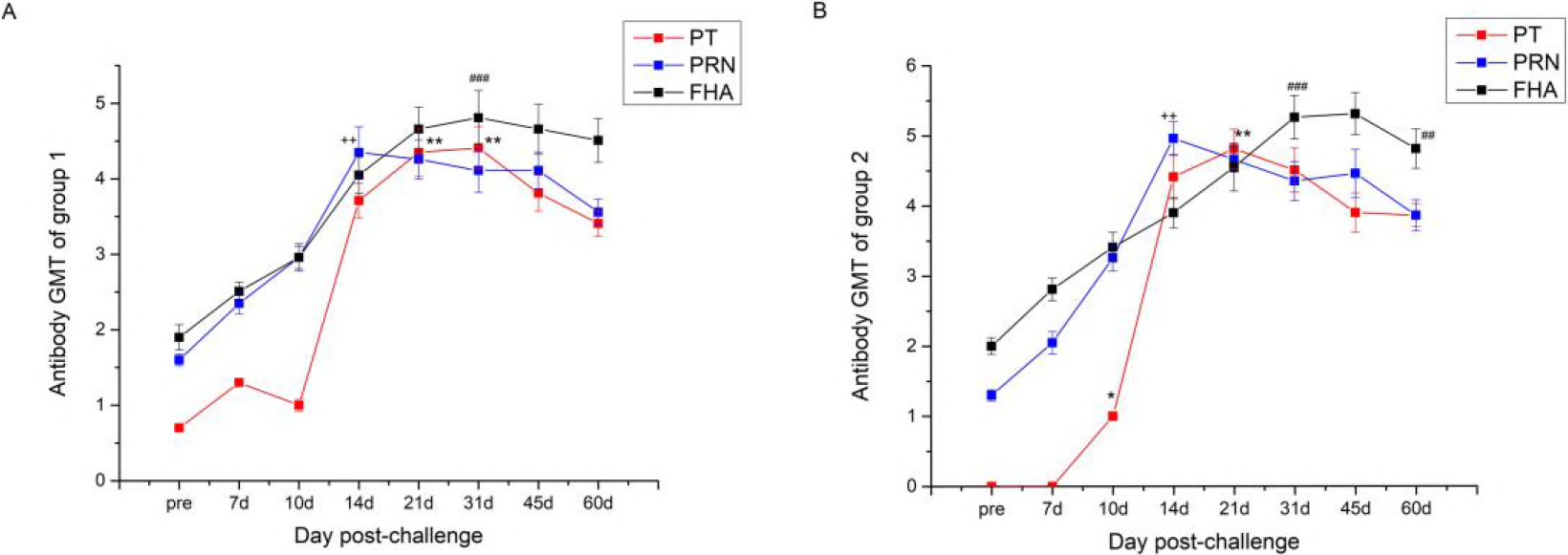
Dynamic profiles of the anti-PT, RPN, FHA antibody titers after challenged by *B.Pertussis*. ELISA method was adopted to determine antibody titers. (A) GMT of anti-PT, RPN, FHA antibody in serum of group challenged for 30 min, *p<0.05, **p<0.01, ++p<0.01, ###p <0.001. (B) GMT of anti-PT, RPN, FHA antibody in serum of group challenged for 60 min, *p<0. 05, **p<0.01, ++p<0.01, ###p<0.001.

### 2.5 Aerosol challenge with *B. pertussis* induced dynamic cytokine responses

The level of sixteen cytokines—GM-CSF, G-CSF, IFN-γ, IL-2, IL-10, IL-15, IL-17A, IL-13, IL-1β, IL-4, IL-5, IL-6, TNFα, IL-12 (p40), IL-18, and IL-8—was measured at each time point after challenge. The results indicated that IL-10 and IL-6 in the serum at day 5 after challenge were significantly increased in group 1 (*P*<0.05) (Fig. 7). In group 2, IL-8 increased significantly at day 5 post-challenge, IL-6 increased significantly at day 7 post-challenge (*P*<0.05), and IL-2, IL-17A, IL-12 (p40), and IL-18 significantly increased at day 14 and decreased at day 21 post-challenge. (*P*<0.05) (Fig. 8). IL-10 increased significantly from day 14 to day 21, and decreased at day 31. IL-13 increased from day 10 and reached the highest at day 14, then decrease but still significant hither than pre-post, finally decreased at day 31.

**Fig. 7.**
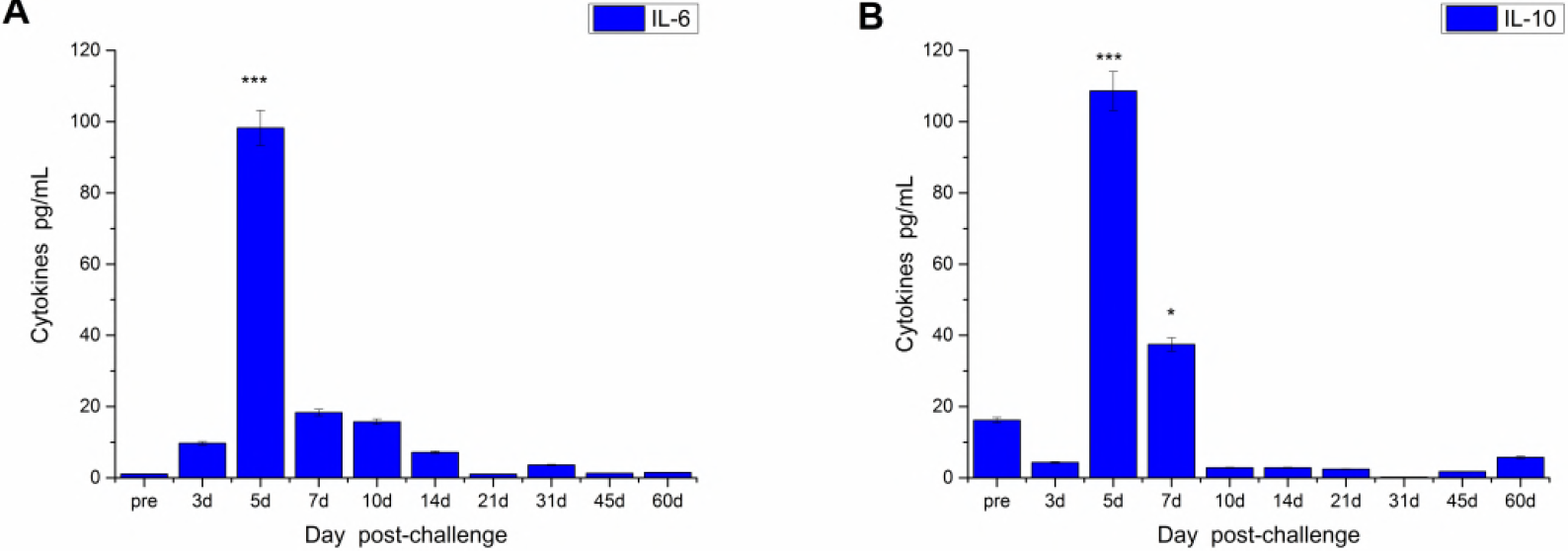
Dynamic profiles of cytokines after challenged 30min with *B. Pertussis*. Venous blood of two groups was drawn at each time point after aerosol challenging and cytokines were determined by liquid chip method. *p<0.05, ***p<0.001.

**Fig. 8.**
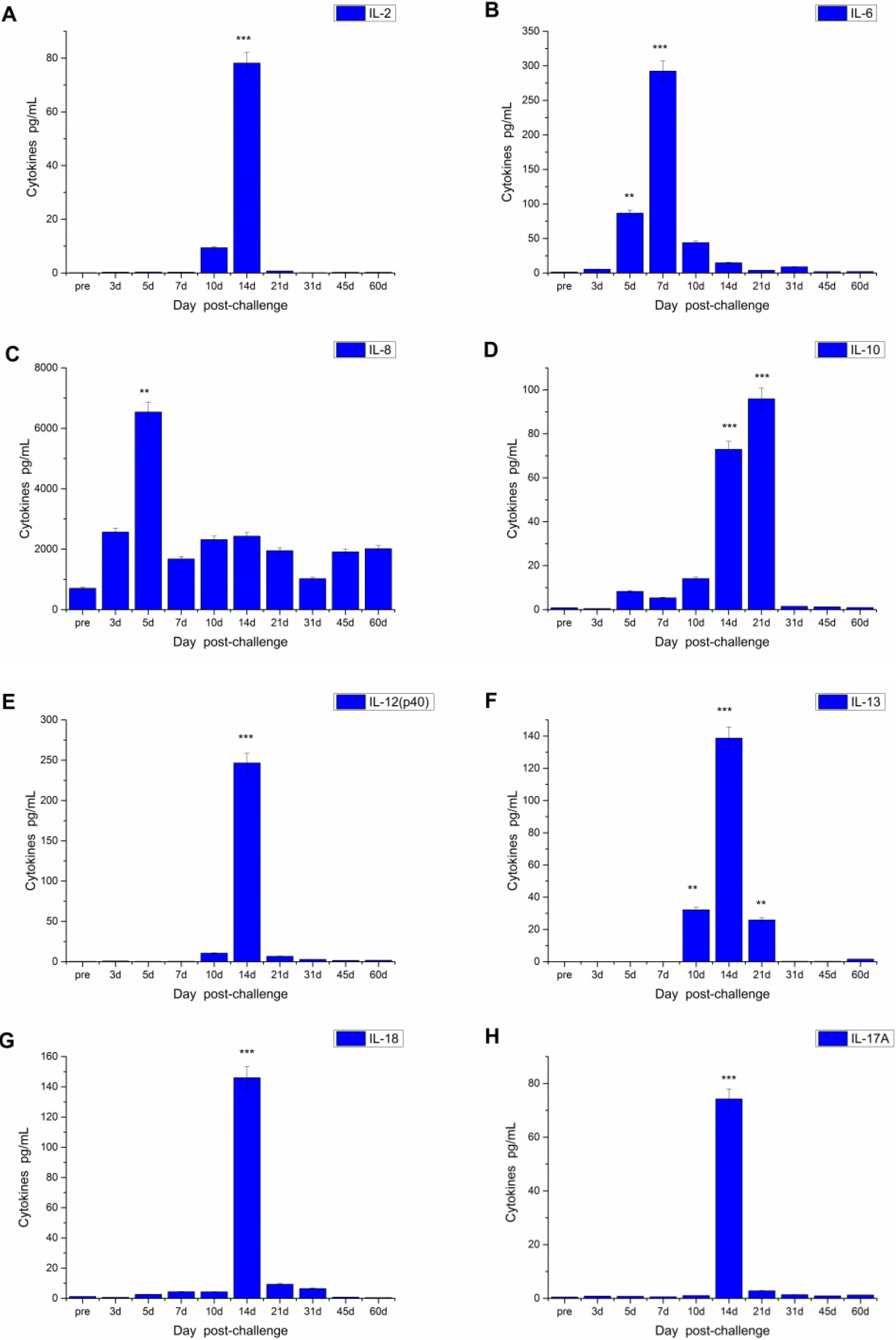
Dynamic profiles of cytokines after challenged 60min with *B. Pertussis*. Concentrations of cytokines in serum at different time point after challenge for 60 min were determined by liquid chip method using Luminex 200 which was described in Fig 7.

## 3 Discussion

Whole-cell pertussis vaccine (wP vaccine) was developed more than 80 years ago. This vaccine significantly reduced the incidence of pertussis in children. However, because of strong adverse reactions, at present most developed countries use acellular pertussis (aP) vaccine. The aP vaccine is produced by different manufacturers and contains different types and amount of pertussis antigens, including pertussis toxin (PT), pertactin (PRN), filamentous haemagglutinin (FHA) and fimbriae (FIM). Moreover, the different immunization schedules for these vaccines increase the variability in vaccine immunogenicity. Some studies indicated that the resurgence of pertussis in recent years might be related to the lower efficacy of aP vaccines[29, 30]. Other studies suggested that the evolution of *B. pertussis* under the pressure of mass vaccination might induce immune adaptation[20]. Therefore, the establishment of suitable experimental challenge models, particularly nonhuman primate models, may provide a platform for studying pertussis pathogenesis, effectiveness of pertussis vaccine, and evaluate the different effectiveness of pertussis vaccine made from different antigen components.

Animal models of infection should reproduce the whole clinical spectrum of pertussis. Compared to other animal models, models with nonhuman primates are ideal because of their close evolutionary relationship with humans. Moreover, these models can help understand infection and the development of pathogenesis[31]. In the present study, rhesus monkeys were selected to investigate the whole clinical spectrum of pertussis after challenge, including inspiratory whoop, posttussive emesis, leukocytosis, and decreasing pulmonary capacity. One of the most important factors to consider when studying the above systems is the method of challenge. To date, nasal challenge, endotracheal intubation, in vivo injection, and aerosol challenge have been investigated in mice models of pertussis[32, 33]. Compared to other challenge methods, aerosol challenge can accurately simulate natural infections and decrease animal stress[34]. In our study, rhesus macaques were infected with *B. pertussis* via aerosol challenge and developed overt signs of clinical pertussis. First, after challenge with 10^5^ CFU/mL *B. pertussis*, dynamic changes in leukocytosis were analyzed. The number of WBCs was significantly increased at day 3, peaked at day 7, and decreased gradually after that, and returned to baseline levels at day 45. The post-challenge concentration of WBCs was 6-fold higher than the baseline level. Second, the number of colonies in nasopharyngeal swabs peaked at day 14 after challenge, reached 10^6.73-6.91^ per mL and returned to baseline levels after 31 days. Bacterial sequencing indicated that colonies were present only in nasopharyngeal samples but not in nasal samples. Third, the rectal temperature of rhesus macaques was increased at days 3 to 7 after challenge, and coughing was observed. However, lacking of suitable video recording system, the coughing frequency and duration were not recorded in present study. In the following research, these should be investigated in detail to provide the further information.

Another critical factor to consider when establishing an animal model is the age of the animals. A previous study using an enterovirus type 71 (EV71) rhesus monkey model suggested that, of the challenged animals, a clinical spectrum similar to that of humans was observed only in young animals[35]. Previous studies that challenged rhesus monkeys with pertussis investigated the fold-change in WBC counts and cough but did not evaluate clinical signs at adults animal[26, 28]. Huang et al [36]has changed Taiwan monkey (Macaca cyclopsis) using 18-323 of H. pertussis and have investigated that the whooping cough, dynamic changes in leukocytosis and antibody response but not fever at young monkey. However it just mentioned the weighing of monkeys was from 550 to 1875 gm, but not the age information be recorded. A study that used baboons as a successful non-human primate pertussis model reported that young baboons showed severe disease signs whereas adult baboons showed mild signs[31]. In the present study, rhesus monkeys aged 12 to 14 months were selected. In our study, the body temperature of rhesus macaques after challenge was increased 1~1.5 °C, which is similar to the increase observed in baboons infected with *B. pertussis* by the endotracheal route[24]. In the baboon model, the peak WBC count was 5~10-fold higher than baseline counts in nine baboons. In our study, the peak WBC count was 4.5~8.4-fold higher than the baseline count. *B. pertussis* was found in nasopharyngeal samples of rhesus macaque. In addition, the bacterial count reached 8.1 × 10^6^/mL and was gradually decreased, which is similar to the results obtained using the baboon model. Therefore, we deduced that the change route as well as the age of the animals may influence the infection of B pertussis.

Except for clinical symptoms, humoral and cellular immune responses are essential to evaluate pertussis infection. Previous studies that investigated immune mechanisms suggested that human pertussis-specific immune response protects against disease rather than against infection[37, 38]. In the present study, anti-PT, PRN, FHA antibody titers were gradually increased 10 days after challenge, anti-PT antibody titers peaked at days 21 to 31, anti-PRN antibody titers peaked at day 14, and anti-FHA antibody titers peaked at days 31 to 45 and then decreased. The specific T cell response to *B. pertussis* associated with different cytokines also plays an important role in the disease. The analysis of T cell responses in children demonstrated that Th1-type responses were predominant in natural infections and wP vaccine injection, whereas Th2-type responses were predominant after aP vaccine injection[39, 40]. Th17 was also involved in protective immunity against *B. pertussis.* Antigen-specific IL-17 was detected in the lungs 7 days post challenge and reached a peak 3 to 4 weeks post challenge[41]. In present study, we measured changes in cellular activity in the serum of the study animals after challenge and found that IL-10 and IL-6 were significantly increased at day 5 in group 1 whereas IL-10, IL-2, IL-17A, IL-13, IL-12 (p40), IL-18, and IL-8 were increased at day 14 in group 2. This result demonstrated that Th1 and Th2 response was induced by aerosolised *B. pertussis,* particularly in group 2. Th17 responses, which are critical for elimination of pertussis, were also detected.

The WBC count, number of colonies in nasopharyngeal samples, and anti-PT, PRN, and FHA antibody titers were significantly higher in group 2 compared to group 1. The levels of IL-10 and IL-6 in group 1 were significantly increased after challenge whereas more cytokines were increased in group 2, indicating that group 2 produced stronger cellular and humoral immune responses and presented more obvious symptoms compared to group 1. These observations could be relevant to high infection doses. The Th1, Th2, and Th17 responses observed in group 2 were similar to those found in the baboon model. Our results demonstrated that aerosol-challenged rhesus macaques aged 12 months could serve as an animal model of pertussis, and a bacterial concentration of 10^5^ CFU/mL for 60 min was a suitable challenge intervention.

The type of challenge bacteria is another critical factor of study of animal model for pertussis. The baboon animal model conducted by Warfel et al [24] used B. pertussis strain D420, a recent clinical isolate from human infant with severe respiratory distress. In present study, CMCC58030(18323), which identified by Pearl Kendrick in the 1940s[42] was used. This stains also been used to challenge Taiwan Monkey in Huang *et al*’s study and showed the experimental whooping cough sighs. The recent analysis on global population structure of B. pertussis has suggested that the prevalence of B. pertussis has changed in the last 100 years worldwide, in which strain 18323 belongs to the branch contains of a small number of stains showing a long distance from the major prevalent branch[43]. However strain 18323 is still used for determination of potency for vaccines in China[44]. Surely, the prevalent stain isolated recently could provide a more powerful information, and it should be used to monitor pathogenicity difference of animal response in the future study.

In summary, the characteristic of pertussis infection in infant rhesus macaque was similar as in human beings, which provide a clue to using infant rhesus macaque as a candidate model of pertussis infection in the future studies for analyzing pertussis infection mechanisms and pertussis vaccines.

## Acknowledgments

We are grateful to the research staff at the Department of Nonhuman Primate Research Center of the Institute of Medical Biology, Chinese Academy of Medical Sciences & Peking Union Medical College for their contribution to this research, and to the Department of Diphtheria, Tetanus, and Pertussis vaccine and Toxins of the National Institutes for Food and Drug Control for assisting in antibody detection.

## Author Contributions

Conceived and designed the experiments: Mingbo Sun, Li Shi. Performed and experiments: Dachao Mou, Peng Luo, Jiangli Liang, Qiuyan Ji, Lichan Wang, Na Gao, Qin Gu, Chen Wei, Yan Ma. Data Analysis: Peng Luo, Dachao Mou, Jingyan Li and Shuyuan Liu. Manuscript Writing: Mingbo Sun, Li Shi, Jingyan Li and Dachao Mou.

